# Centromeres evolve progressively through selection at the kinetochore interface

**DOI:** 10.1101/2025.01.16.633479

**Authors:** Jana Helsen, Kausthubh Ramachandran, Gavin Sherlock, Gautam Dey

**Affiliations:** Department of Genetics, Stanford University School of Medicine, Stanford, CA, USA; Cell Biology and Biophysics, European Molecular Biology Laboratory, Heidelberg, Germany; Collaboration for joint PhD degree between EMBL and Heidelberg University, Faculty of Biosciences, Heidelberg, Germany

## Abstract

During mitosis, stable but dynamic interactions between the centromere DNA and kinetochore complex enable accurate and efficient chromosome segregation. Even though many proteins of the kinetochore are highly conserved, centromeres are among the fastest evolving regions within a genome, showing extensive variation even on short evolutionary timescales. Here, we sought to understand how new types of centromeres emerge and reach fixation by mapping centromere evolution across 138 budding yeast species and over 2,500 natural strain isolates. We show that new centromeres spread progressively via drift and subsequent selection, and that the kinetochore interface, which is evolving slowly in relative terms, determines which new centromere variants are tolerated. Together, our findings provide insight into the evolutionary constraints and trajectories shaping centromere evolution.

## Introduction

Centromeres are indispensable chromosomal elements for cell division across eukaryotes. As attachment points for the chromosome segregation machinery, they are responsible for partitioning the cell’s DNA rapidly and reproducibly through a stable but dynamic interaction with the kinetochore complex and spindle microtubules (Fig. 1A). Although each centromere needs to accomplish this essential cellular feat alongside a conserved set of kinetochore proteins *(1, 2),* centromeres are among the fastest evolving regions in the genome (*3, 4*), showing striking variability across the tree of life (*5, 6*). Centromeres range from complex epigenetically defined regions of hundreds of kilobases embedded in megabase-sized arrays of satellite DNA in metazoans (*7*) and plants (*8*), to genetically defined loci of 100-200 nucleotides in budding yeasts (*9*).

**Figure 1.**
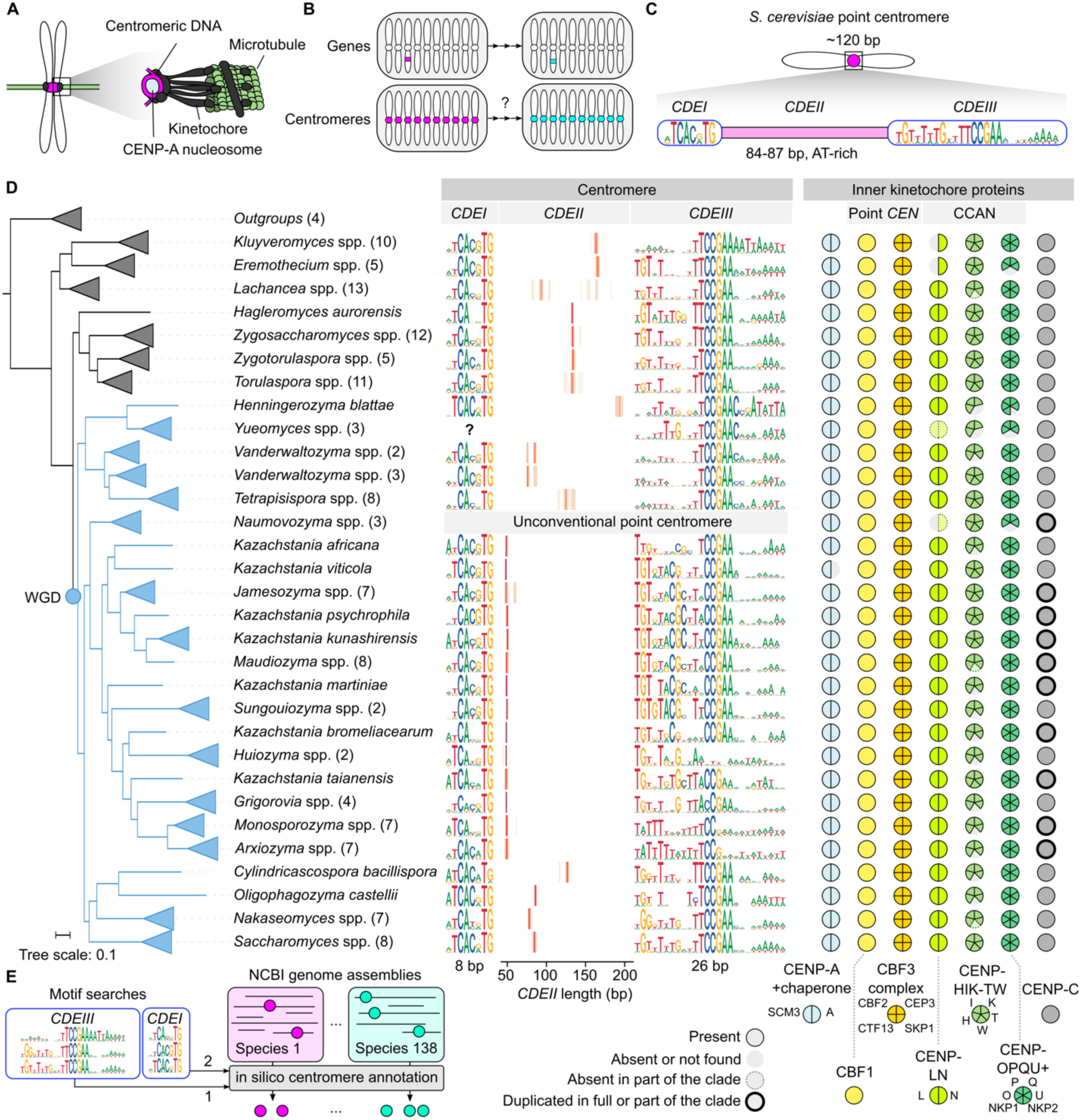
The point centromere sequence landscape. (**A**) Stylized schematic of centromeric DNA bound to a microtubule through the kinetochore complex. (**B**) Schematic highlighting the complexity of centromere evolution. (**C**) The structure of the *Saccharomyces cerevisiae* point centromere. *S. cerevisiae* point centromeres are defined by an ∼85 bp-long AT-rich region (*CDEII*) flanked by two DNA motifs (*CDEI* and *CDEIII*). (**D**) Landscape of predicted centromere sequences across 138 *Saccharomycetaceae* and accompanying absence/presence profiles of inner kinetochore proteins. Species phylogeny was determined using a concatenation-based maximum likelihood analysis of 1,270 orthologous groups of proteins under a single LG+G4 model. Branches belonging to the same genus are collapsed and are represented by triangles. The numbers of species belonging to each collapsed clade are indicated in parentheses next to the genus names. Branches of species that emerged after the whole genome duplication (WGD) are colored in blue. Centromere sequences are represented by DNA logos for *CDEI* and *CDEIII*, with graphs indicating *CDEII* length in between. *Naumovozyma* spp. have unconventional point centromeres lacking regular *CDEI* and *CDEIII* motifs (*17*). Centromere profiles and predictions for *Naumovozyma* spp. can be found in Fig. S2 and Data S2. Point *CEN*: point centromere-specific kinetochore proteins. CCAN: constitutive centromere-associated network. (**E**) Simplified schematic representing the in silico point centromere annotation pipeline (PCAn). Centromeres were detected in assemblies through two sequential motif searches: (1) *CDEIII* and (2) *CDEI*.

Unlike protein-coding genes, which are often only present as single copies within the genome, species with monocentric chromosomes possess multiple centromeres, one on each chromosome. For organisms to evolve centromeres with novel features, this means that each chromosome must either alter its existing centromere or acquire a novel centromere (Fig. 1B), making centromere evolution conceptually very different from gene evolution. To date, many of the basic evolutionary principles underlying centromere evolution remain largely unknown. It remains unclear (i) whether centromere transitions occur progressively or concurrently, (ii) how much they are a result of selection and/or drift, and (iii) how these rapidly evolving regions ensure that their crucial connections with the kinetochore and spindle are maintained. In this study, we use a combination of centromere discovery, phylogenetic profiling, and *in vivo* centromere function experiments to determine the evolutionary constraints and trajectories of centromere transitions in budding yeasts.

## Results

### Mapping the point centromere sequence landscape

To systematically explore the evolutionary mechanisms driving centromere transitions, centromere diversity needs to be mapped across the phylogenetic tree in the context of recent evolution. We selected the Saccharomycetaceae, a fungal clade including the model species *Saccharomyces cerevisiae* and *Nakaseomyces glabratus* (formerly *Candida glabrata*) to identify and characterize such transitions as they have short, genetically defined ‘point’ centromeres (*10*), use a structurally similar spindle to segregate their chromosomes (*11*), and the clade encompasses many species within a relatively short evolutionary time frame (*12*). Each point centromere is defined by an AT-rich region (*CDEII*) and two DNA motifs (*CDEI* and *CDEIII*) (Fig. 1C) (*13*) that are bound by specific DNA-binding proteins (Fig. 1D), several of which are unique to the clade (*14*). While it is known that point centromeres vary across the clade, sequences are only available for a handful of species (*9, 14*), insufficient to pinpoint when and how centromere transitions occurred. To compile a systematic list of centromere sequences across the *Saccharomycetaceae* clade, we developed an automated <u>P</u>oint <u>C</u>entromere <u>An</u>notation tool (PCAn) which uses two sequential motif searches to detect point centromeres in genome assemblies (Fig. 1E) (see Material and Methods for details).

Using PCAn to annotate centromeres for 138 species (Data S1), we generated a comprehensive atlas of point centromere diversity (Fig. 1D, Fig. S1, Fig. S2, Data S2). The numbers of predicted centromeres correspond well with published chromosome numbers (Fig. S3), validating our approach while highlighting that our tool can also be used to obtain karyotype information (Fig. S1). Despite the limited diversity in inner kinetochore composition, point centromeres show extensive diversity across the clade. In some lineages, parts of the *CDEIII* motif are either completely conserved or eroded away (e.g., *Huiozyma* spp. lack the conserved CCGAA motif, while part of the *CDEIII* motif is more conserved in *Sungouiozyma* spp.). However, perhaps the most striking variation can be observed in the length of *CDEII*, the AT-rich region that wraps around the centromeric nucleosome (*15, 16*). *CDEII* length ranges from around 50 base pairs in many *Kazachstania*-related species to nearly 200 base pairs in *Henningerozyma blattae*. Notably, while there is significant *CDEII* length variation between species, the length distribution within any given genome is remarkably consistent (Fig. S1). Our atlas of centromere diversity shows that point centromeres have evolved extensively since their origination, and that centromere transitions occurred multiple times within the *Saccharomycetaceae* clade.

### Centromeres transition progressively through selection and drift

Next, we used our comprehensive atlas of centromere diversity to explore the evolutionary dynamics underlying centromere transitions. Given the substantial interspecific variation yet limited intraspecific variability in *CDEII* length, we focused on the evolutionary trajectories of this key centromere feature. Full transitions of *CDEII* length occurred independently on multiple occasions (Fig. 2A-C), even within the same genus (Fig. 2B-C). Unexpectedly, we also found a few species carrying two distinct *CDEII* lengths simultaneously, i.e., species in which two distinct centromere types coexist within the same genome (Fig. 2C-D). This suggests these species are currently in a state of centromere transition. Within the *Jamesozyma* genus, while most species retain the ancestral ∼50 bp *CDEII*, *J. spencerorum* has undergone a complete transition to a ∼60 bp *CDEII*, whereas its sister species, *J. jinghongensis*, has a nearly equal distribution of both centromere types (Fig. 2C). Another example of mixed centromere states is found in the *Vanderwaltozyma* genus. All species within this genus possess a mixture of centromeres with either a ∼75 bp or ∼85 bp *CDEII*, with the relative frequency of each variant varying across the clade. While some species exhibit an equal distribution of both centromere types, others are significantly enriched for either the shorter or longer variant (Fig. 2D). These observations indicate that centromere transitions occur progressively: centromeres change one by one, with a transitional phase characterized by the coexistence of old and new centromere variants within the same genome.

**Figure 2.**
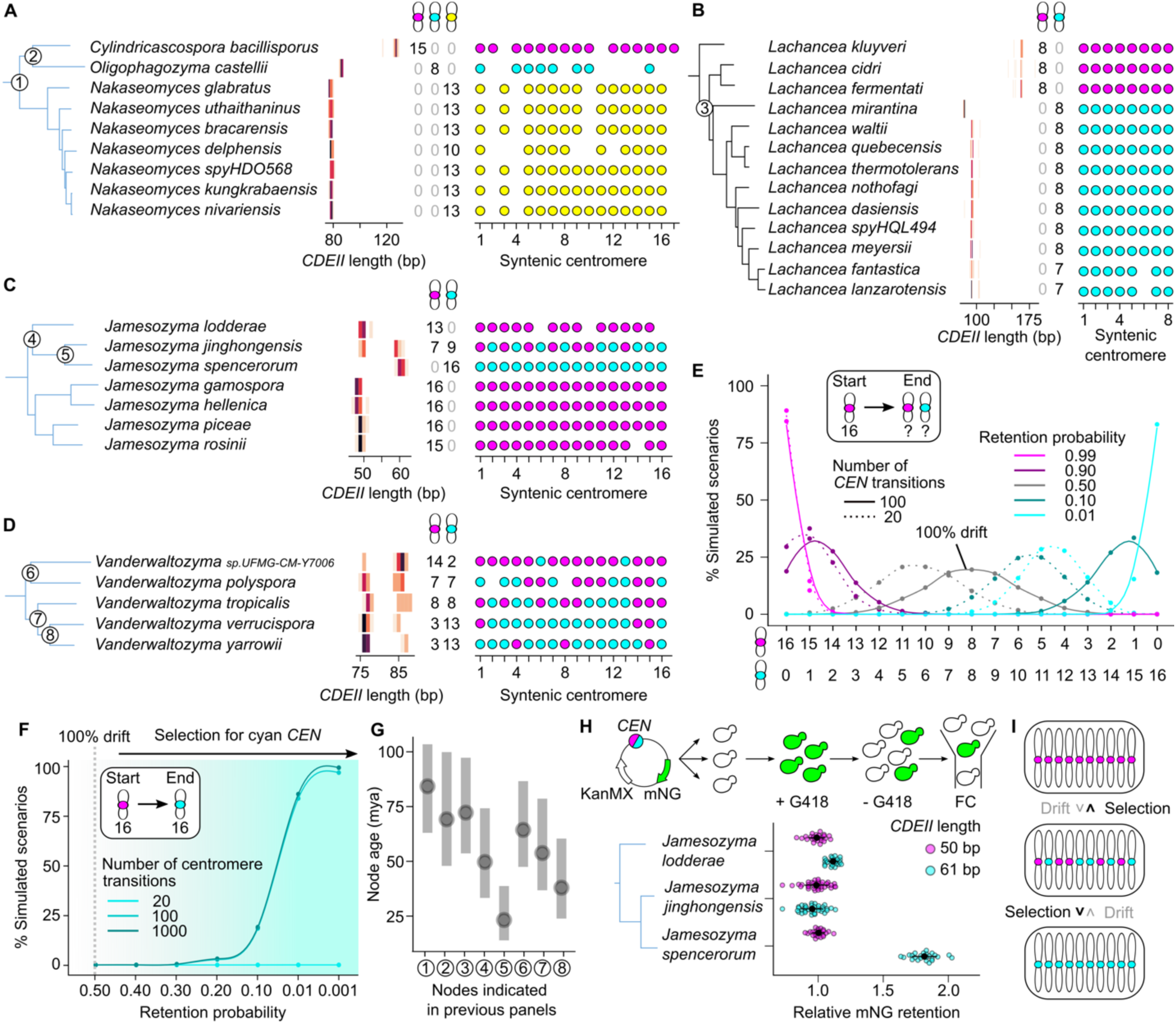
Centromeres transition gradually through drift and selection. (**A-D**) *CDEII* length across species within the (**A**) *Cylindricascospora, Oligophagozyma,* and *Nakaseomyces* genus, (**B**) *Lachancea* genus, (**C**) *Jamesozyma* genus, and (**D**) *Vanderwaltozyma* genus. Magenta, cyan and yellow indicate different centromere types, as defined by different *CDEII* lengths. The total number of each centromere type and their syntenic locations are indicated on the right. The numbered nodes represent full or partial centromere transition points. (**E**) Simulated centromere transitions. Starting from 16 magenta centromeres, each transition, one random centromere was drawn and transitioned from magenta to cyan or cyan to magenta. The chance that this new variant is retained was then determined by the retention probability, where a 100% retention implies that magenta variants will always be retained (i.e., selection for magenta), a 50% retention rate implies that there is an equal chance the new variant is retained or lost (i.e., drift), and a 0% retention rate implies that magenta variants are never retained (i.e., selection for cyan). This was done 20 (dotted lines) or 100 times (full lines), and the simulation was repeated 10,000 times. The x-axis gives all possible scenarios at the end of the simulation (i.e., the number of magenta and cyan centromeres), and the y-axis gives the proportion of the 10,000 simulations for each of these scenarios. (**F**) Simulated full centromere transitions. The simulations were done the same way as described for panel E but specifically focused on the scenario of full transitions (16 magenta centromeres to 16 cyan centromeres). The x-axis represents different retention probabilities and the y-axis gives the proportion of the 10,000 simulations in which we observe a full transition for seven different retention probabilities. There were no simulations in which full transitions are achieved without selection. (**G**) Node age (mya: million years ago) of the nodes indicated in panels A-D. (**H**) Plasmid loss assay in three *Jamesozyma* species. A centromeric plasmid with selectable marker (KanMX: conferring resistance to G418) and fluorescent protein (mNG: mNeonGreen) was transformed into *J. lodderae, J. jinghongensis*, and *J. spencerorum*. To measure the mitotic efficiency of different centromere variants, cells were first grown in selective medium (+G418) and were then grown for a set number of generations in non-selective medium (-G418) after which the proportion of fluorescent cells was determined using flow cytometry (FC). The graph gives the relative retention rate of a plasmid carrying a centromere with a 50 bp *CDEII* (magenta) or 61 bp *CDEII* (cyan) for each species. (**I**) Schematic of proposed evolutionary model for centromere transitions. *n =* 24.

Next, we sought to determine the role of selection in these progressive transitions. If different centromere variants can coexist in one cell, this implies that both can establish a sufficiently stable connection with the mitotic machinery. However, some variants might still be better at segregating and will thus be favored by selection. To determine if selection is required to achieve full centromere transitions, we conducted *in silico* centromere transition experiments (Fig. 2E-F, Fig. S4). These simulations show that full centromere transitions are indeed not possible without selection for the new variant (Fig. 2E-F). Even in a scenario where the ancestral and new variants are equally good at segregating (drift alone causing frequency changes), full transitions are impossible. Similarly, selection for ancestral centromeres is required to maintain a full complement of ancestral variants (Fig. 2E, Fig. S4). In the examples shown in Fig. 2A-D, such full transitions took between ∼23 and ∼84 million years of evolution (Fig. 2F). To determine whether different centromere variants do indeed show differences in segregation efficiency before and after full transitions, we determined the in vivo retention rate of plasmids carrying centromeres with either a 50 bp or 61 bp *CDEII* in three *Jamesozyma* species (Fig. 2G). Only *J. spencerorum*, the species that underwent a complete transition to centromeres with a ∼60 bp *CDEII*, has a strong preference for the longer variant, indicating that selection was indeed required for the centromere transition in this clade. We hypothesize that centromere transitions occur progressively through a combination of selection and drift (Fig. 2I): differences in segregation efficiency determine which variants will be favored by selection, and which new variants are neutral and can spread through drift.

### Centromere variants spread through populations through sex

While the data shown above demonstrate the progressive nature of centromere transitions, they do not inform us about the mechanisms underlying the initial emergence and subsequent spread of novel centromere variants within populations. To address this, we focused on intraspecific variation. Employing PCAn to annotate centromeres in 1,493 unique *S. cerevisiae strains* (Data S3) revealed that, while the majority contain 16 centromeres with consistent *CDEII* lengths (∼85 bp), approximately 9.5% harbor one or two variants with significantly deviating *CDEII* lengths (< 80 bp or > 90 bp) (Fig. 3A, Fig. S5, Data S4). To determine whether these 142 strains all carry the same variant centromere or 142 different ones, or, in other words, how many times independent variants arose and spread in the population, we assigned synteny and aligned centromere sequences and were able to assign 13 distinct variant centromeres within the *S. cerevisiae* population (Fig. 3B). Notably, some variants are found across different clades and in mixed/mosaic clades (Fig. 3B-C), indicating that centromere variants can spread across populations through sex. This is further supported by the observation that strains can carry a combination of two different variants, and that some centromere variants are present heterozygously (Fig. 3D). By including sexual cycles in our simulations, we show that full centromere transitions can spread through populations more easily (Fig. 3E). Finally, we investigated the underlying mutational mechanisms driving the emergence of these novel centromere variants. Remarkably, the majority of variant centromeres seem to have expanded through a microhomology-mediated mutational mechanism (Fig. 3F). This is not only true for centromere variants in *S. cerevisiae*, but also for variants detected in other *Saccharomycetaceae* (Fig. S6). One hypothesis is that these small homologous insertions are the result of stalled replication forks (*18*), which is a frequent occurrence at budding yeast centromeres (*19*).

**Figure 3.**
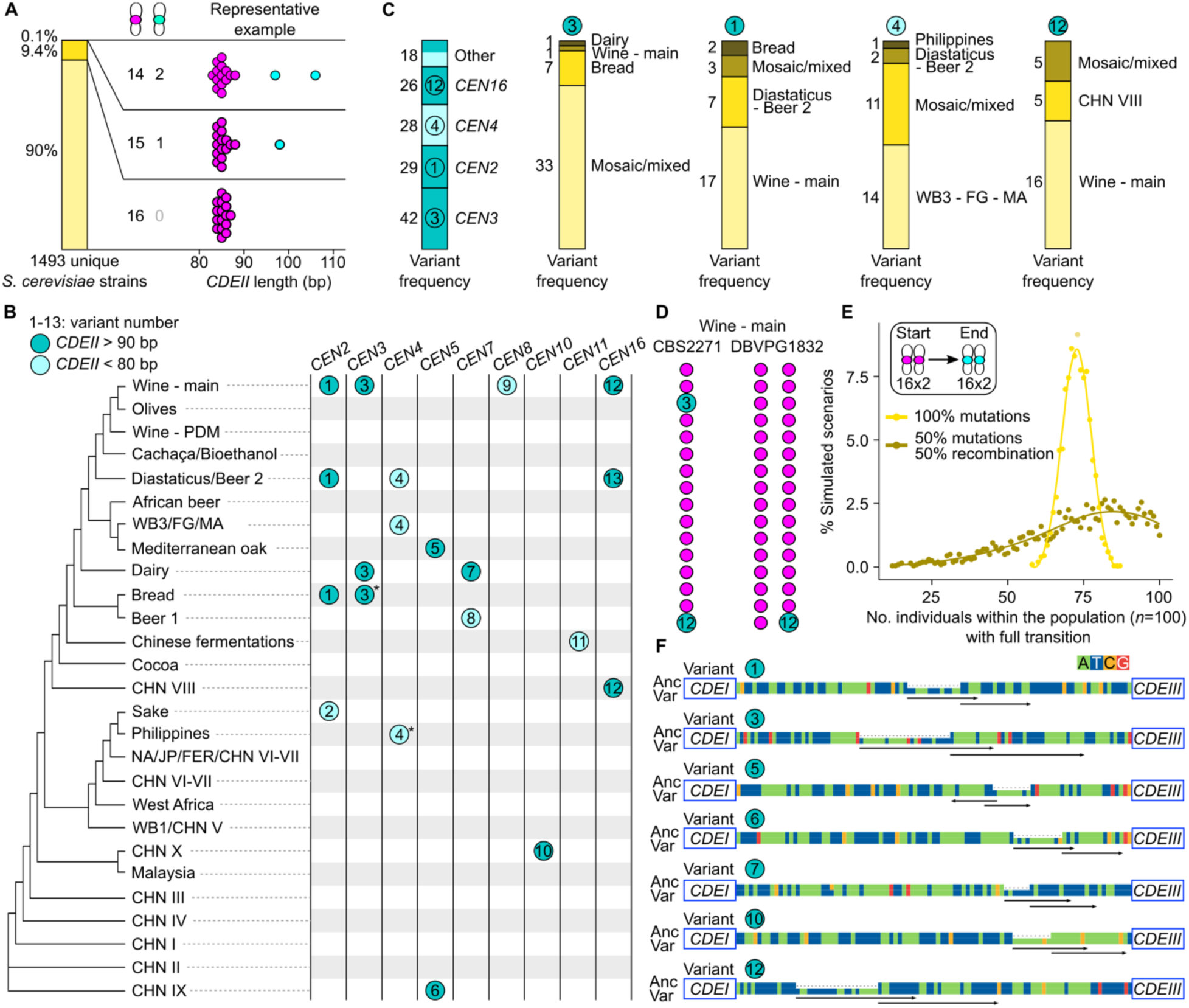
Centromere variants spread through populations through sex. (**A**) Proportion of *Saccharomyces cerevisiae* strains with ‘regular’ centromeres (magenta, 80-90 bp *CDEII*) and variant centromeres (cyan). (**B**) Spread of variant centromeres throughout the natural *S. cerevisiae* population. Light blue circles represent variants with *CDEII* < 80 bp and dark blue circles represent variants with *CDEII* > 90 bp. The numbers in the circles represent independent variants. Centromere IDs are indicated on top and correspond to chromosome IDs in *S. cerevisiae* (e.g., *CEN2* is the centromere on ChrII). Clades and tree structure were taken from (*20*). (**C**) Prevalence of the most frequent variants across different clades. (**D**) Example of a *S. cerevisiae* strain with two variant centromeres (left) and a strain in which the variant centromere is heterozygous (right). (**E**) Simulated full centromere transitions in populations with diploids with and without recombination. Starting from 16x2 magenta centromeres in 100 individuals, each step, one random individual was drawn in which one random centromere was drawn and transitioned from magenta to cyan or cyan to magenta. The retention probability of the magenta variant was 1%. In the condition with recombination, this step was followed by randomly drawing new sets centromeres for each chromosome of each individual in the population, based on the prevalence of each variant type within the population. The simulation was repeated 2,000 times, with 500 steps in each simulation. The x-axis represents the number of individuals within the population with full transitions, and the y-axis gives the proportion of the 2,000 simulations in which we observe a full transition. (**F**) Mutations underlying *CDEII* length variation in *S. cerevisiae*. Variant numbers correspond to the numbers indicated in the previous panels. Variant sequences (Var) were aligned with the most similar ‘ancestral’ sequences (Anc). Black arrows indicate identical sequences.

### Centromere transitions are constrained by the kinetochore interface

Can centromeres transition to just any new state, or are there constraints on the types of changes that can occur? While the centromere atlas in Fig. 1 shows that point centromeres, and *CDEII* length in particular, can vary significantly over longer evolutionary timescales, the examples shown in Fig. 2 and Fig. 3 suggest that transitions can only occur through a select set of mutations. In both examples, most mutations lead to a ∼10 base pair jump in *CDEII* length. The ability to tolerate such jumps in centromere length seems to be a defining feature of the point centromere clade (Fig. 4A). As nucleosomal DNA twists at ∼10.2 base pairs per turn (*21*), we hypothesize that a ∼10 base pair jump in *CDEII* length ensures that the *CDEI* motif stays at the orientation necessary for Cbf1 to properly bind the motif and interact with other components of the kinetochore complex (Fig. 4B). Indeed, by measuring the segregation efficiency of centromeres with different *CDEII* lengths in *S. cerevisiae*, we found that variants that are ∼10 base pairs longer than wild-type centromeres show no segregation defects while centromere variants that are only 5 base pairs longer are lost more readily (Fig. 4C). Together, these observations indicate that the kinetochore interface dictates which centromere variants can be tolerated and subsequently spread through populations.

**Figure 4.**
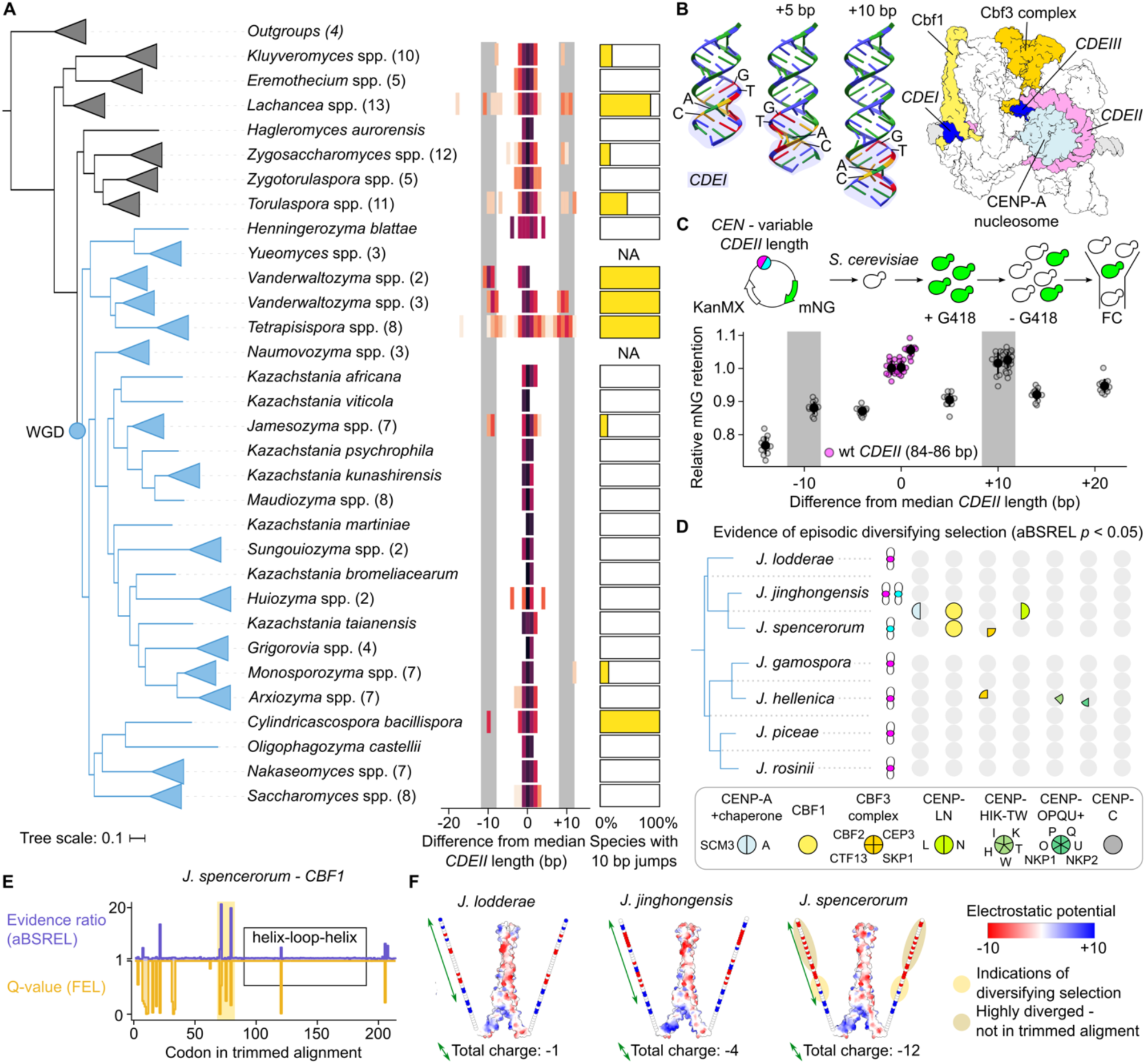
Centromere transitions are constrained by the kinetochore interface. (**A**) *CDEII* length variation profile across 138 Saccharomycetaceae, represented by the difference from the median *CDEII* length and the proportion of species with 10 base pair *CDEII* length variation for each clade. Branches belonging to the same genus are collapsed and are represented by triangles. The numbers of species belonging to each collapsed clade are indicated in parentheses next to the genus names. Branches of species that emerged after the whole genome duplication (WGD) are colored in blue. (**B**) Relative orientation of the *CDEI* motif on pieces of DNA of different length and the structure of the inner kinetochore in *Saccharomyces cerevisiae* highlighting the DNA motifs (dark blue), AT-rich region (*CDEII*: pink) and motif-binding proteins (Cbf1: yellow and Cbf3 complex: orange). Other inner kinetochore proteins are colored in white and pieces of DNA not part of *CDEI, CDEII* or *CDEIII* are colored in grey. Structure from (*22*). (**C**) Plasmid loss assay with different *CDEII* length in *S. cerevisiae*. Centromeric plasmids with varying *CDEII* length, a selectable marker (KanMX: conferring resistance to G418) and fluorescent protein (mNG: mNeonGreen) were transformed into *S. cerevisiae*. To measure the mitotic efficiency of different centromere variants, cells were first grown in selective medium (+G418) and were then grown for a set number of generations in non-selective medium (-G418) after which the proportion of fluorescent cells was determined using flow cytometry (FC). The graph gives the relative retention rate of a plasmid carrying centromeres with different *CDEII* lengths. The most prevalent wild-type lengths are indicated in pink. *n* = 12. (**D**) Internal and external branches in the *Jamesozyma* genus with evidence of episodic diversifying selection (aBSREL *p* < 0.05) for different inner kinetochore proteins. Chromosomes with magenta or cyan centromeres indicate centromere type in each species. (**E**) Evidence ratio (aBSREL) and Q-value (contrast-FEL) across the trimmed alignment of *CBF1* in *J. spencerorum*. The yellow box highlights a region enriched for mutations indicative of diversifying selection. (**F**) Structures and electrostatic potential of Cbf1 dimers for *J. lodderae, J. jinghongensis* and *J. spencerorum*. Structures were predicted using Alphafold2, and only segments with pLDTT > 70 are shown as volumes. The region indicated in panel E falls in the N-terminal tail, a part of which is represented by circles. Positively charged residues are represented by blue circles and negatively charged residues are represented by red circles.

Finally, while this accounts for how new neutral centromere variants can spread through drift, this does not explain how new variants might become more efficient and favored by selection, as observed in *Jamesozyma* spp. in Fig. 2H. To explore if this might be the result of adaptation of the kinetochore machinery itself, we tested the inner kinetochore proteins in the genus for evidence of positive selection. Coincident with centromere transitions, Cbf1, the protein that binds the *CDEI* motif (Fig. 4B), shows evidence of episodic diversifying selection (Fig. 4D). Most of the signal comes from a small segment of the protein’s N-terminal tail, close to the bHLH leucine zipper domain which binds *CDEI* (Fig. 4E) (*22*). Cbf1’s N-terminal tail is a fast-evolving disordered region whose removal only leads to a minor reduction in segregation efficiency in *S. cerevisiae* (*23*). In *J. spencerorum*, the species that recently underwent a complete centromere transition and in which the new centromere variant became the more efficient variant, the mutations resulted in a new acidic patch close to the zipper domain (Fig. 4F, Fig. S7), potentially influencing Cbf1’s overall orientation through interactions with other proteins of the kinetochore. These findings suggest that coevolution between the kinetochore and the evolving centromere sequences could play a role in driving centromere transitions.

## Discussion

With our new tool for point centromere annotation, we expanded the limited repertoire of available tools for clade-specific *in silico* centromere prediction (*24–26*) and generated a clade-wide overview of point centromere diversity. We showed that centromeres transition progressively through a combination of drift, selection, and sex, to new states that are compatible with the kinetochore interface. We propose a model where the initial fate of a new centromere variant is largely determined by its compatibility with the existing kinetochore machinery. If the variant can establish a sufficiently stable interaction with the kinetochore, it can spread within the population through neutral processes such as drift and sexual reproduction. Subsequent changes in the segregation efficiency of different variants, either through modifications to the kinetochore machinery (e.g., mutations in kinetochore proteins like Cbf1) or through environmental changes (e.g., temperature changes impacting microtubule dynamics) could then create selective pressures that ultimately drive full centromere transitions. Indeed, several kinetochore proteins show signatures of positive selection, some of which correlate with evolving centromeric features (*27–32*). One popular model used to explain rapid centromere evolution is the centromere drive model, which proposes that the most efficient centromere variants are preferentially passed on to the next generation during asymmetric female meiosis (*33*). However, this model may not fully explain centromere evolution in the Saccharomycetaceae, as many species in this clade strictly undergo male meiosis where all four meiotic products are viable. Additionally, our results show that differences in segregation efficiency during mitosis, too, can vary significantly between species. In unicellular organisms, both sexual (∼germline) and asexual (∼somatic) variation are passed on to the next generation. Our work underscores that centromere evolution is in fact the result of an interplay of various factors, including drift and selection during both mitosis and meiosis, following evolutionary trajectories that are constrained by the kinetochore interface.

## Material and Methods

### Genomes

Genome assemblies for each of the 138 *Saccharomycetaceae* and four outgroup species (*Wickerhamomyces anomalus, Candida albicans, Pichia kudriavzevii,* and *Yarrowia lipolytica*) were downloaded from NCBI (Data S1). To explore *S. cerevisiae* intraspecies centromere variation, 2,737 assemblies were downloaded from NCBI and from the supplemental data of four studies (*34–37*) (Data S3). As some strain backgrounds (e.g., S288c) are overrepresented in this dataset, we limited our selection to 1,493 unique strains with phylogenetic information for Fig. 3. Regardless, the overall frequency of variant centromeres was very similar between the reduced and full dataset (Fig. S5).

### Point centromere annotation pipeline (PCAn)

#### PCAn pipeline

FIMO (*38*) from the MEME suite (version 4.11.2) (*39*) was used to search each genome assembly with a chosen 26 bp-long *CDEIII* motif (see below), using a threshold ranging from 1.0e-3 to 1.0e-7. The coordinates of the hits were used to compile a list of 250 bp-long sequences that contain the *CDEIII* motif plus 224 bp upstream. Subsequently, this output was searched again using FIMO and a chosen 8 bp-long *CDEI* motif with a less restrictive threshold of 1.0e-2. The coordinates of these hits were used to compile a list of sequences that contain a *CDEI* motif on one end and a *CDEIII* motif at the other end. For each of these sequences, the length and AT% of the intermediary *CDEII* motif were calculated. Every sequence was assigned a combined score based on the two FIMO hit scores and the *CDEII* AT%. Sequences were sorted based on the combined score and only the top 50 sequences were retained. Next, we determined the median *CDEII* length of the top 5 sequences and removed sequences with lengths differing more than 30 nucleotides from the median. Finally, after removing sequences with a low *CDEII* AT content (<70%) and removing duplicates (sometimes found on small contigs of lower-quality assemblies), we removed sequences in which both *CDEII* length and AT content differ too much from the median (≷ median ± 10 nucleotides *and* AT% - 7, respectively).

#### Motif construction and selection

*CDEI* and *CDEIII* sequences from (*9*) were used to produce the initial search motifs, using the ‘sites2meme’ command from the MEME suite (version 4.11.2) (*39*). Newly discovered sequences were each verified using synteny and used to make new motifs and perform more sensitive searches within specific clades or species. The final PCAn pipeline uses the best-performing custom motifs and thresholds for each clade or species.

#### Synteny checks

Synteny checks were performed by identifying the proteins 10 kb up- and downstream of centromere hits. After using contig and coordinate information of each centromere hit to extract the ∼20 kb region around each potential centromere, open reading frames (ORFs) were identified using the getORFProteins function from ORFFinder Python (version 1.8) (minimum_length = 525, remove_nested = True, return_loci = True) (*40*). ORFs were then BLASTed against the *S. cerevisiae* proteome using a local version of NCBI’s blastp (version 2.13.0+, default parameters) (*41*). Finally, the *S. cerevisiae* protein identifiers were matched to the ancestral *Saccharomycetaceae* gene order identifiers from the Yeast Gene Order Browser (*42*) which were used for visualization.

#### Inner kinetochore protein searches

Based on a literature review and the list used in (*22*), we selected 21 inner kinetochore proteins in *S. cerevisiae*. We used tblastn (BLAST suite version 2.5.0+, default parameters) (*41*) to retrieve the coordinates for the ORFs in the genomes of the 137 other Saccharomycetaceae. We rejected all hits with an E-value greater than 1e-10. For the remaining hits, we then used the getORFProteins option (minimum_length=210, remove_nested=True, return_loci=True) in ORFFinder Python (version 1.8) (*40*) to identify all possible ORFs within 5 kb on either side of the starting coordinate. We selected the ORFs which contained the midpoint of the selected 10kb scaffold. We further selected only those ORFs which, on performing blastp against the *S. cerevisiae* proteome, returned the correct starting protein as the best hit. Finally, for each instance where this approach failed to retrieve homologs, we manually checked and confirmed the presence/absence of homologs using phylum-specific homology searches.

### Species tree construction

#### Selecting marker proteins

We inferred the species tree for the set of 142 species (138 *Saccharomycetaceae* and four outgroup species: *Wickerhamomyces anomalus, Candida albicans, Pichia kudriavzevii,* and *Yarrowia lipolytica*) using 1,403 marker genes identified and published in an earlier study (*43*). From this set of 1,403 markers, 113 markers were removed as they did not have a corresponding protein homolog in *S. cerevisiae*. We mapped the markers to the *S. cerevisiae* reference proteome using phmmer from HMMER suite of tools (*44*) and removed 13 markers that returned the same best hit as other markers in the dataset, resulting in a set of 1,277 markers.

#### Retrieving homologs for marker proteins

We constructed BLAST databases using makeblastdb (BLAST suite version 2.5.0+) (*41*) for each of the 142 genomes. We used tblastn (default parameters) to retrieve the ORF coordinates for the 1,277 markers from the 142 genomes. We used the getORFProteins option in ORFFinder Python (version 1.8) (*40*) to identify the entire ORF for each potential homolog with an E-value < 1e-10 and only selected those homologs which returned the corresponding marker protein in the *S. cerevisiae* proteome as the best hit in a reverse blastp search. At this stage, we excluded 7 more markers for which we were only able to identify reciprocal best hits in less than 50% of the selected 142 species.

#### Constructing gene trees, removing outliers, and reconstructing and dating the species tree

For each of the 1,270 sets of homologs, we used MAFFT v7.505 (*45*) with the E-INS-i option to align the sequences, trimal v1.4.rev15 build[2013-12-17] (*46*) with the “-gappyout” option to remove phylogenetically noisy positions, and FastTree Version 2.1.11 Double precision (No SSE3) (*47*) with “-spr 4 -mlacc 2 -slownni -n 1000 -gamma” options to build maximum-likelihood (ML) gene trees. We analysed these trees using ETE3 (*48*) in Python to identify and remove branches in the tree with branch lengths greater than 20 times the median branch length. In cases where the median branch length was less than 1e-8, we manually inspected the trees and alignments to remove outliers. These steps were repeated until no outliers were found.

We concatenated the 1,270 alignments obtained after outlier removal into one supermatrix alignment with 672,500 positions using Goalign (*49*). We used IQ-TREE multicore version 2.2.0.3 (*50*) with “-B 1000 -alrt 1000 --boot-trees --wbtl -m LG+G4 -mwopt --threads-max 24 -T AUTO” options to build a ML tree using the LG model (*51*) with 4 rate categories (LG+G4) with 1,000 ultrafast bootstraps (*52*). We used the LG+G4 model as it was the selected model for 661/1,270 of the marker genes.

To estimate divergence times in a tractable manner, we randomly subsampled the supermatrix alignment and extracted 3 sets of 10,000 sites and inferred ML trees using IQ-TREE as described earlier. We inferred the time tree for these three trees using the RelTime-ML method (*53*) implemented in MEGA11 (*54*). We adopted two well-estimated ranges of divergence from two internodes as calibration points: the *Saccharomyces cerevisiae* – *Saccharomyces uvarum* split (14.3 mya – 17.94 mya) and the *Saccharomyces cerevisiae* - *Kluyveromyces lactis* split (103 mya – 126 mya) (*12, 55*).

### Centromere transition simulations

#### Haploids without recombination

Each experiment was initialized with one individual carrying 16 centromeres of type ‘A’. Next, numpy’s random number generator (seed: 19680801) was used to generate a random number from 0 to 15 to pick one random centromere. After this, a second random number between 0 and 999 was generated and compared to the retention probability to decide if the chosen centromere will be forced to transition or not. The retention probability always refers to how likely it is to retain the original type ‘A’. For example, if the chosen centromere is of type ‘A’ and the retention probability is 0.99, numbers from 0 to 989 will lead to retention of type ‘A’, and numbers from 990 to 999 will lead to a transition to type ‘B’. Similarly, if the chosen centromere were of type ‘B’, numbers from 0 to 989 will lead to transition to type ‘A’, and numbers from 990 to 999 will lead to retention of type ‘B’. This process was repeated 20, 100 or 1,000 times for different retention probabilities. The simulations were repeated 10,000 times.

#### Diploids with and without recombination

Each experiment was initialized with one hundred individuals carrying 2x16 centromeres of type ‘A’. Each iteration was composed of a mutation step, similar to what is described above for haploids, followed by a recombination step for the condition with recombination. For the mutation step, numpy (seed: 19680801) was used to pick a random individual in the population, then a random chromosome pair, and then a random chromatid. Then, exactly the same as for haploids, another random number was generated and compared to the retention probability to decide if the chosen centromere will be forced to transition or not. For the condition with recombination, each individual in the population was then allowed to undergo ‘meiosis’: for each chromosome, a random chromatid was retained and the other chromatid was randomly picked from the population. This process was repeated 500 times, with and without recombination. The simulations were repeated 2,000 times.

### Plasmid loss assays

To measure the relative retention rates of plasmids containing different centromere variants, we used the same principle as employed in (*56*). Apart from components necessary for propagation in *E. coli*, each plasmid contained an autonomously replicating sequence shown to function across different *Saccharomycetaceae* (*57*), a *KANMX6* cassette for selection in yeast, a *pADH1-mNeonGreen-tADH1* construct for constitutive expression of mNeonGreen, and a variable centromere sequence. Plasmid sequences can be found on the FigShare repository accompanying this manuscript. Plasmids were transformed into *J. lodderae* (CBS 2757), *J. jinghongensis* (CBS 15232), *J. spencerorum* (DBVPG 6746) and *S. cerevisiae* (BY4741), using the standard LiAc-based transformation protocol for budding yeast.

For the loss assays, cells were grown overnight in 10 mL YPAD supplemented with G418 disulfate (Carl Roth): 800 µg/mL for *Jamesozyma* spp. and 400 µg/mL for *S. cerevisiae*. Except for *J. jinghongensis*, which was grown at 25°C, all other species were grown at 30°C throughout the experiment. The next morning, the OD_600_ was measured, cultures were washed in YPAD and diluted to OD_600_ = 1 in YPAD. A sample was taken to determine the proportion of fluorescent cells at T0. The cultures were then diluted 1:256 in YPAD (200 µL in 50 mL) and 150 µL of these cultures was pipetted into wells of a 96-well plate. Cells were grown for 24h, after which the proportion of fluorescent cells was measured by flow cytometry on an Accuri C6 (BD), or a FACSymphony A3 (BD) (minimum 30,000 cells).

### Protein evolutionary analyses

For every *Jamesozyma* spp. and each of the inner kinetochore proteins identified above, we extracted the corresponding gene sequence using the same combination of tblastn (BLAST suite version 2.5.0+, default parameters) (*41*) and ORFFinder Python (version 1.8) (*40*). Protein sequences were aligned using MAFFT v7.505 (*45*) (default parameters) and protein alignments were then converted to codon alignments using PAL2NAL v14 (*58*) (default parameters). Gaps and ambiguously aligned sites were removed using Gblocks v0.91b (*59*), using -t = c; -b1 = $b; - b2 = $b; -b3 = 1; -b4 = 6; -b5 = h, with $b the number of sequences divided by two plus one, as applied in (*60, 61*). aBSREL v2.5 (*62*) and contrast-FEL v0.5 (*63*) from the HyPhy suite (using default parameters) were then used to look for signatures for episodic diversifying selection and differences in selective pressure at individual sites in *J. spencerorum*, respectively.

### Alphafold2-multimer modeling of Cbf1 dimers

We predicted the structures for the Cbf1 dimers from *J. lodderae, J. jinghongensis* and *J. spencerorum* using a local installation of Colabfold 1.5.5 (*64*) (--num-recycle 12 --num-ensemble 1 --model-type auto --save-pair-representations). We used the entire Cbf1 sequence as retrieved from the homology searches. Colabfold used MMseqs2 (*65*) on specific, clustered databases (*66, 67*) to retrieve homologs for Cbf1 and construct alignments. AlphaFold2-multimer (*68*) with 12 recycles was used to predict the structure of the Cbf1 dimer. The resulting structure predictions were visualized using UCSF ChimeraX v1.8 (*69*).

## Data and code availability

PCAn and all other code related to this project can be found on GitHub: https://github.com/JHelsen/point-centromere-detection

Supplementary data accompanying this manuscript can be found on FigShare: https://doi.org/10.6084/m9.figshare.c.7630151.v2

## Acknowledgements

We thank Dr. Tom A. Williams from the University of Bristol for his invaluable discussions and suggestions with regards to the species tree reconstruction, and Dr. Michelle Hays for invaluable critical feedback on the manuscript. We are also grateful for the resources and assistance of the EMBL IT services and HPC cluster (https://doi.org/10.5281/zenodo.12785829). Funding was provided by the Life Science Alliance (Bridging Excellence Fellowship) (J.H.); NIGMS R35 GM131824 (G.S.); European Molecular Biology Laboratory (J.H., K.R. and G.D.); the European Union (ERC, KaryodynEvo, 101078291) (J.H. and G.D.), and the Joachim Herz Stiftung (Add-on Fellowship for Interdisciplinary Life Science) (K.R.).

## Declaration of Interests

The authors declare no competing interests.

---

**Fig. S1.**
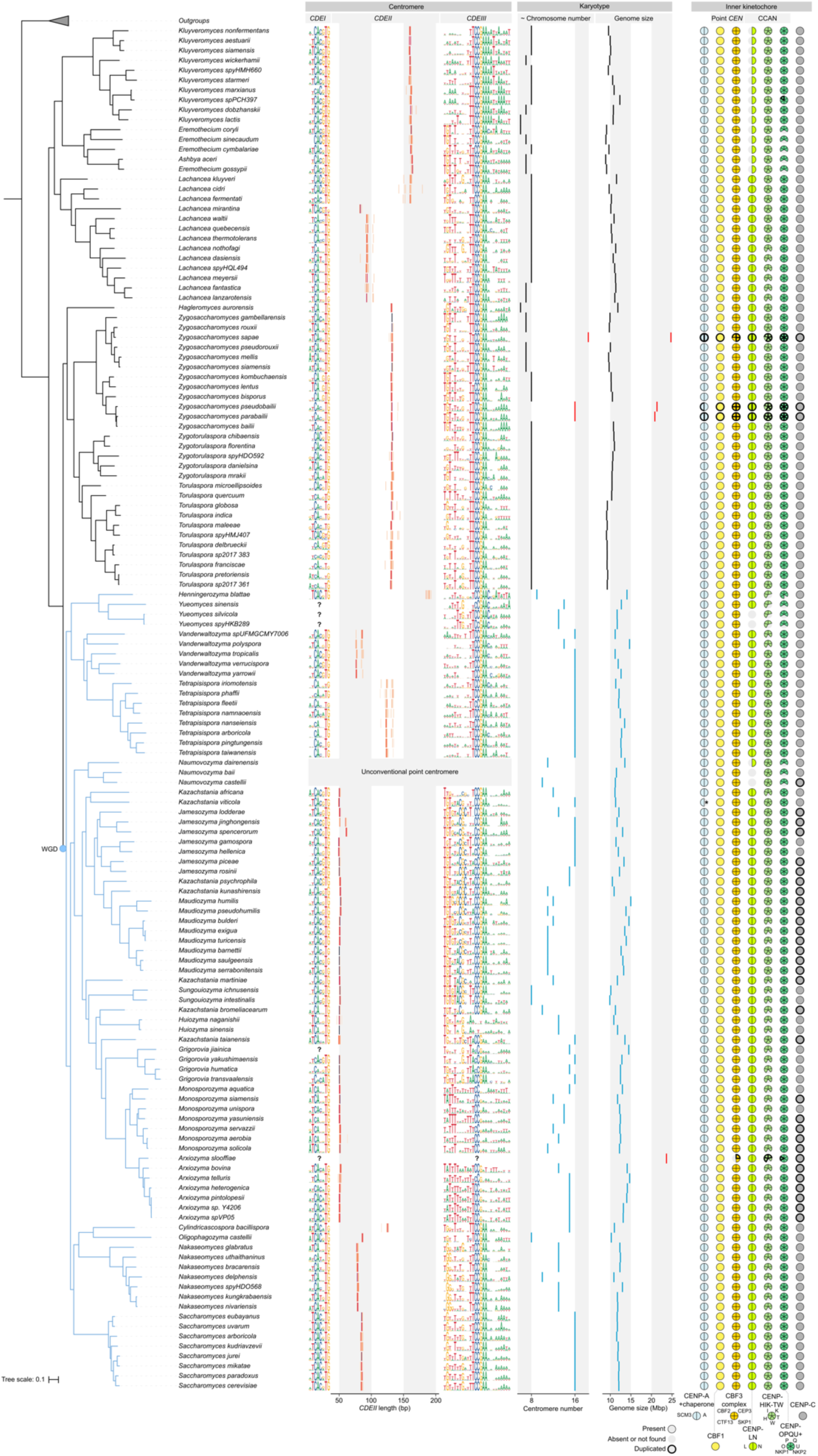
Uncollapsed tree with centromere sequences and inner kinetochore absence/presence protein profiles for 138 Saccharomycetaceae. Species phylogeny was determined using a concatenation-based maximum likelihood analysis of 1,270 orthologous groups of proteins under a single LG+G4 model. Branches of species that emerged after the whole genome duplication (WGD) are colored in blue. Centromere sequences are represented by DNA logos for *CDEI* and *CDEIII*, with graphs indicating *CDEII* length in between. Centromere numbers and genome size for each species are indicated in the middle panels with estimates for hybrid species indicated in red. CCAN: constitutive centromere-associated network. *CENP-A was not detected in *Kazachastania viticola*, but contigs in the genome assembly are broken up in the region where the gene is supposed to be. Hence, we believe its absence is an artifact of the assembly.

**Fig. S2.**
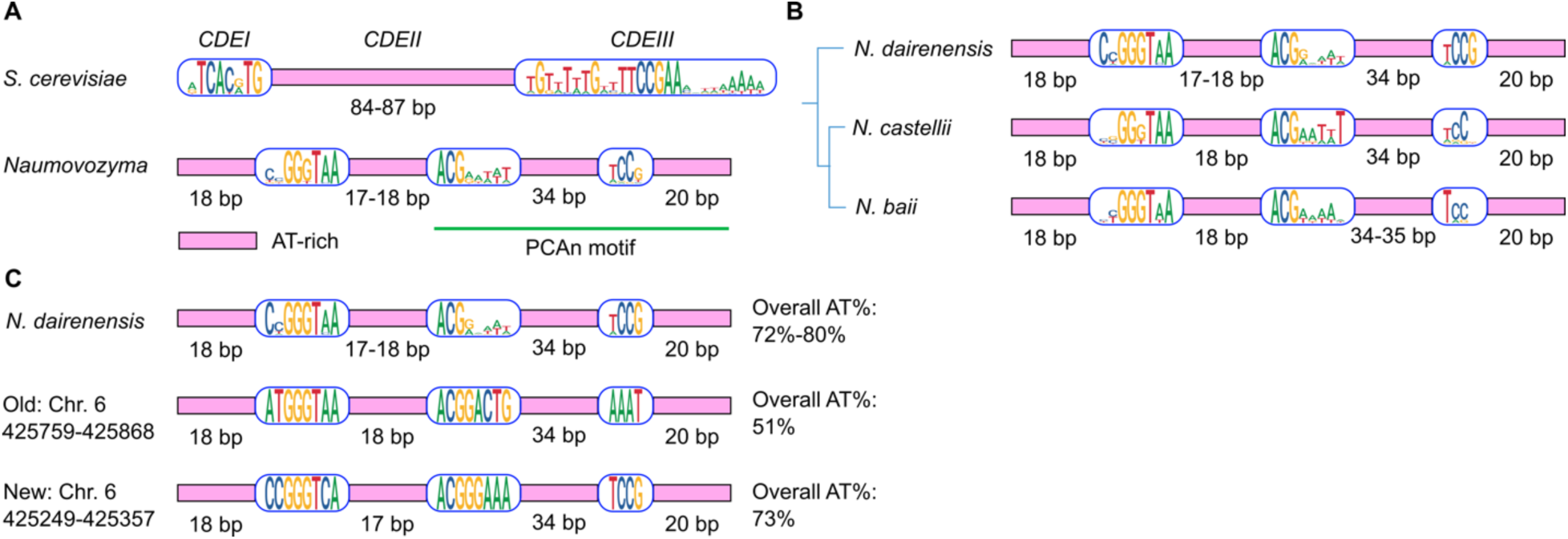
Naumovozyma centromeres. (**A**) *Naumovozyma* spp. lack conventional point centromeres. Instead, they have point centromeres with unique, non-syntenic *CDE*s, indicating that they did not evolve from conventional point centromeres and have an independent evolutionary origin (*17*). To see if our method can be adapted to also identify these types of point centromeres, we adapted PCAn and tried different versions of the motifs identified in (*17*). In contrast to the conventional version of PCAn, what worked best was to use one long 66 bp motif instead of two. (**B**) Using this altered version of PCAn, we were able to correctly identify all ten centromeres in *N. castellii*. In *N. dairenensis*, we identified the correct number of centromeres (eleven), but on chromosome 6 we predicted a sequence different from (*17*) (**C**). Since our hit is very close to the original prediction (and syntenic with the centromere found in *N. castellii*), and a much better match to the consensus motif, we propose that our hit is the ‘real’ centromere sequence of chromosome 6. For *N. baii*, PCAn only managed to detect 6 high-confidence syntenic hits. Either the motif in this species is too different to be picked up by our pipeline, or some centromeres relocated, or the quality of the assembly is insufficient.

**Fig. S3.**
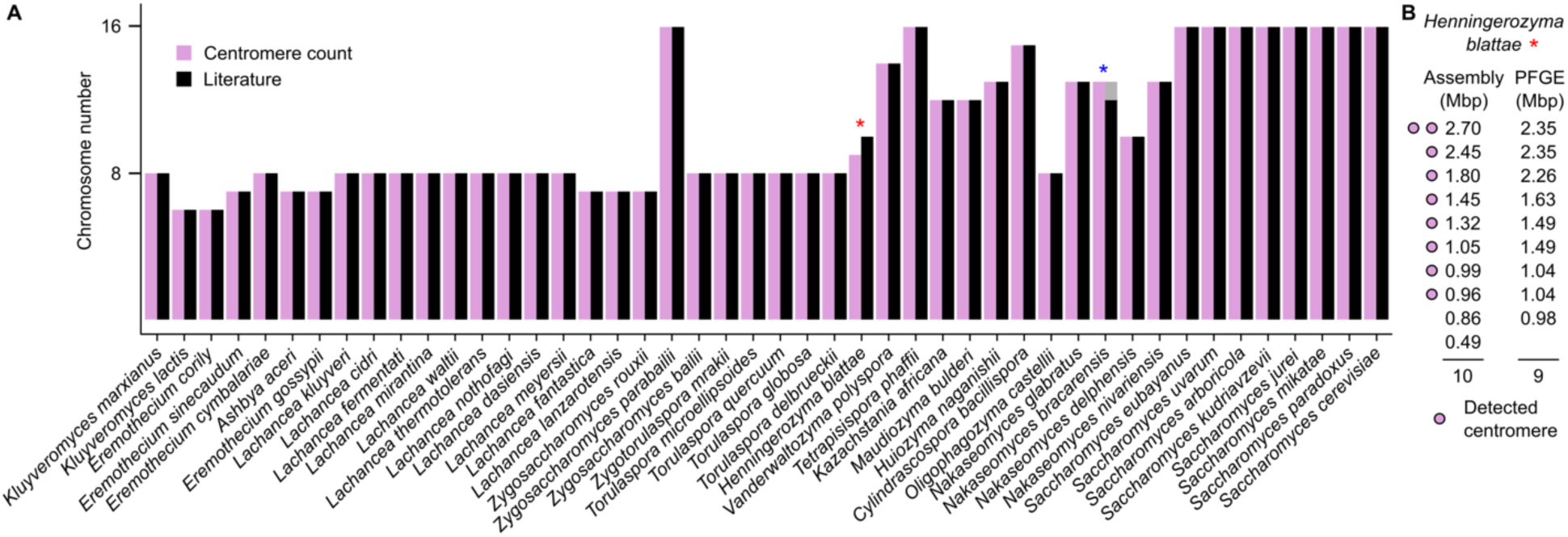
Predicted centromere numbers correspond to chromosome numbers reported in literature. **(A)** Computed centromere counts (plum) versus chromosome numbers reported in literature (black). The blue asterisk indicates a case in which our pipeline identified 13 centromeres and the most recent assembly is unsure whether there are 12 or 13 chromosomes (*70*). The red asterisk indicates one case in which the numbers are not identical. This is further discussed in **(B)**, in which chromosome sizes from the chromosome-level assembly are compared to chromosome sizes observed in PFGE gels of the same strain (*Henningerozyma blattae* CBS 6284) (*71*). Our chromosome number estimate corresponds with the estimate obtained using PFGE gels.

**Fig. S4.**
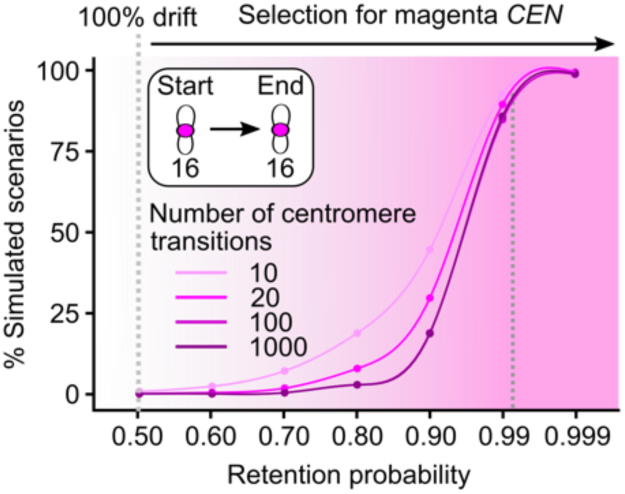
Simulated centromere transitions - full retention. Starting from 16 magenta centromeres, each transition, one random centromere was drawn and transitioned from magenta to cyan or cyan to magenta. The chance that this new variant is retained was then determined by the retention probability, where a 100% retention implies that magenta variants will always be retained (i.e., selection for magenta), a 50% retention rate implies that there is an equal chance the new variant is retained or lost (i.e., drift), and a 0% retention rate implies that magenta variants are never retained (i.e., selection for cyan). Here we specifically focus on the scenario of full retention (16 magenta centromeres remain 16 magenta centromeres). The x-axis represents different retention probabilities and the y-axis gives the proportion of the 10,000 simulations in which we observe a full transition for seven different retention probabilities.

**Fig. S5.**
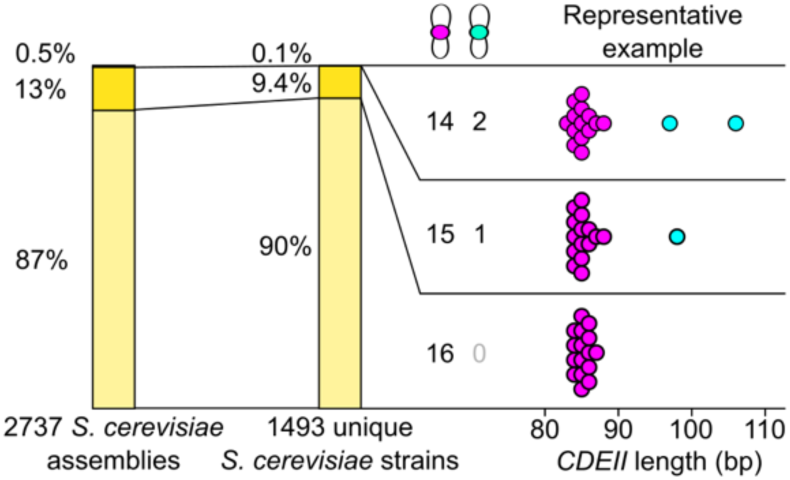
*S. cerevisiae* assemblies and unique strains with variant centromeres. Proportion of *Saccharomyces cerevisiae* assemblies and unique strains with ‘regular’ centromeres (magenta, 80-90 bp *CDEII*) and variant centromeres (cyan).

**Fig. S6.**
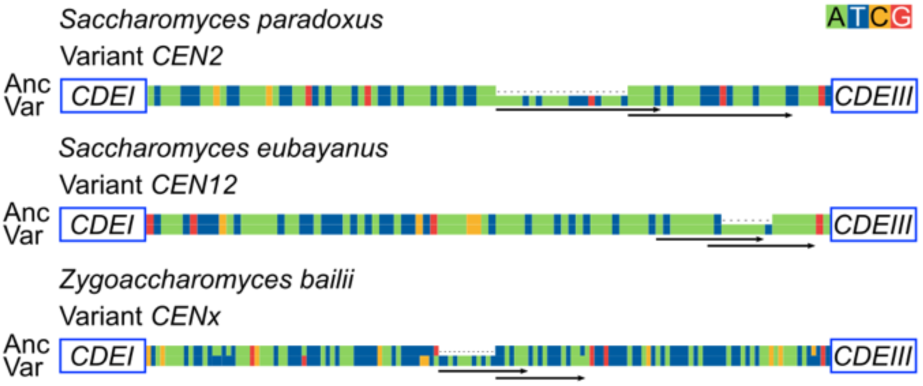
Centromere variants with microhomology in species other than *S. cerevisiae*. Variant numbers correspond to the numbers indicated in the previous panels. Variant sequences (Var) were aligned with the most similar ‘ancestral’ sequences (Anc). Black arrows indicate identical sequences.

**Fig. S7.**
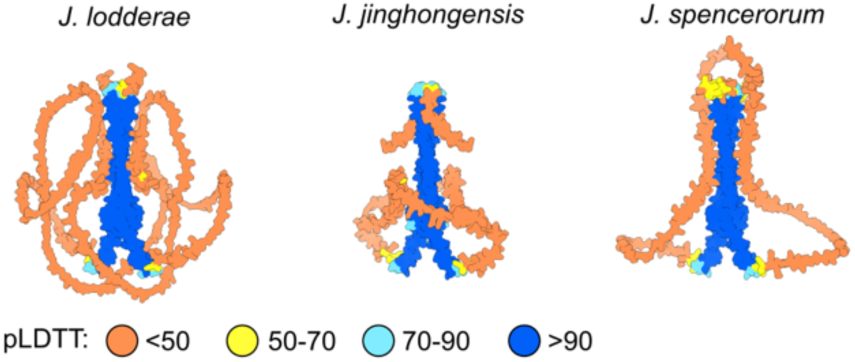
Full Alphafold2 predictions of Cbf1 dimers. Structures of Cbf1 dimers for *J. lodderae, J. jinghongensis* and *J. spencerorum*. Structures were predicted using Alphafold2 and colors represent different pLDTT values.

## Supplemental data

**Data S1 Species and genome identifiers** Species names and GenBank assembly identifiers for species used in this study. Species names correspond to NCBI’s taxonomic classification (January 2025).

**Data S2 Centromere sequences across species** File with every annotated centromere across the 138 Saccharomycetaceae. Columns represent species name, GenBank assembly identifier, genome size, contig, centromere number, full centromere sequence, *CDEI* sequence, *CDEIII* sequence, *CDEII* sequence, *CDEII* length, and *CDEII* AT content.

**Data S3 *S. cerevisiae* strains and genome identifiers** Dataset source and identifiers for the 2,737 *S. cerevisiae* assemblies used in this study. ‘NCBI assemblies’ were downloaded from NCBI, ‘Peter et al. genomes’ can be found in (*34*), ‘Farmhouse genomes’ in (*37*), ‘Almeida genomes’ in (*35*), and ‘Stingless bees’ in (*36*).

**Data S4 Centromere sequences across *S. cerevisiae* strains** Syntenic centromere annotations for all 2,737 *S. cerevisiae* assemblies. The first column indicates the assembly, and the next 32 columns contain syntenic centromere assignments. Column 2_A, for example, contains every annotated centromere on chromosome 2. If there were two centromere variants found for a certain chromosome, the second variant can be found in column_B.

## Notes

https://doi.org/10.6084/m9.figshare.c.7630151.v2

https://github.com/JHelsen/point-centromere-detection

### Competing Interest Statement

The authors have declared no competing interest.

